# Multivoxel patterns for perceptual confidence are associated with false color detection

**DOI:** 10.1101/735084

**Authors:** J.D. Knotts, Aurelio Cortese, Vincent Tascherau-Dumouchel, Mitsuo Kawato, Hakwan Lau

**Author notes:** **Correspondence** (J.K.).

## Abstract

While it has been proposed that metacognition and conscious perception are related, the mechanistic relationship between the two is unclear. To address this question, we combined decoded neurofeedback (DecNef) in functional magnetic resonance imaging (fMRI) with concurrent psychophysics. Participants were rewarded for activating multivoxel patterns for color discrimination confidence while they detected color in mostly achromatic stimuli. We found that occurences of voxel patterns for high color discrimination confidence were associated with false alarms in the concurrent color detection task, suggesting a link between discrimination confidence and consciousness.

## Introduction

Some current theories of consciousness posit a link between consciousness and metacognition (Lau and Rosenthal 2011; Dehaene, Lau, and Kouider 2017). Intuitively, one cannot consciously see something without having some sense of certainty or uncertainty regarding what is being seen (Dienes 2007; Fleming and Lau 2014; Rosenthal 2018); but see (Block 2007). While behavioral evidence supports the idea that metacognitive judgments are a meaningful proxy for conscious experiences (Persaud, McLeod, and Cowey 2007; Dienes and Seth 2010; Szczepanowski et al. 2013; Rausch and Zehetleitner 2016; Norman and Price 2015), the extent of this support has been questioned (Sandberg et al. 2010; Overgaard et al. 2010; Rosenthal 2018; Norman and Price 2015). It has also been suggested that a common mechanism may underlie biases in conscious perception (e.g. conservative detection) and metacognitive misjudgments (e.g. under-confidence in discrimination); in disorders like blindsight, both seem to be problematic (Ko and Lau 2012). And yet, these claims have so far not been directly tested.

We capitalized on the findings of a previous study in which we showed that perceptual confidence could be decoded from multivoxel fMRI patterns in lateral prefrontal and parietal cortex (Cortese et al. 2016). Pairing these patterns with reward modulated participants’ reported confidence in a subsequent dot motion discrimination task. Our question here concerns whether these changes in reported confidence reflect changes in conscious experience too.

To answer this question, we rewarded participants for simultaneously activating decoded voxel patterns for both perceptual confidence in frontoparietal areas (high vs low confidence) and color perception in early visual areas (red vs green stimulus color), while they viewed a stimulus that was achromatic on the majority (> 97%) of trials. During this closed-loop fMRI procedure, we asked participants at regular intervals to report whether they saw any color in the stimulus. We found that when they falsely detected non-existent color, there was an association with occurrences of multivoxel patterns for high color discrimination confidence, supporting the link between a metacognitive process and conscious perception.

## Materials and Methods

### Experiment Overview

The experiment had four main stages across a total of seven days (Figure 1a): the multivoxel pattern analysis (MVPA) sessions (Days 1-2), pre-DecNef psychophysics (Day 3), DecNef (Days 4-6), and post-DecNef psychophysics (Day 7). During the MVPA sessions on Days 1 and 2 participants (N=17) performed a red/green color discrimination task with confidence judgments (Figure 1b) and a color lightness task with both red and green stimuli (Figure 1c) inside an fMRI scanner, and the resulting blood oxygen-level dependent (BOLD) signal patterns were used to train binary decoders for high versus low confidence and red versus green color, respectively. During the DecNef stage participants performed a real-time neurofeedback task in which they were rewarded for activating decoded multivariate BOLD signal patterns corresponding to redness in visual cortex and high confidence in frontoparietal cortex. To examine whether activation of decoded color and confidence patterns had any correspondence with real-time color perception, participants performed a concurrent color detection task during DecNef. Finally, to examine whether this neurofeedback manipulation had any effect on red/green color discrimination (Amano et al. 2016), participants performed the same red/green color discrimination task as in the MVPA sessions outside of the scanner during the pre- and post-DecNef psychophysics stages. Days 2 and 3 always occurred on separate calendar weeks, and were thus always separated by at least two days. Days 3-7 were always consecutive.

**Figure 1.**
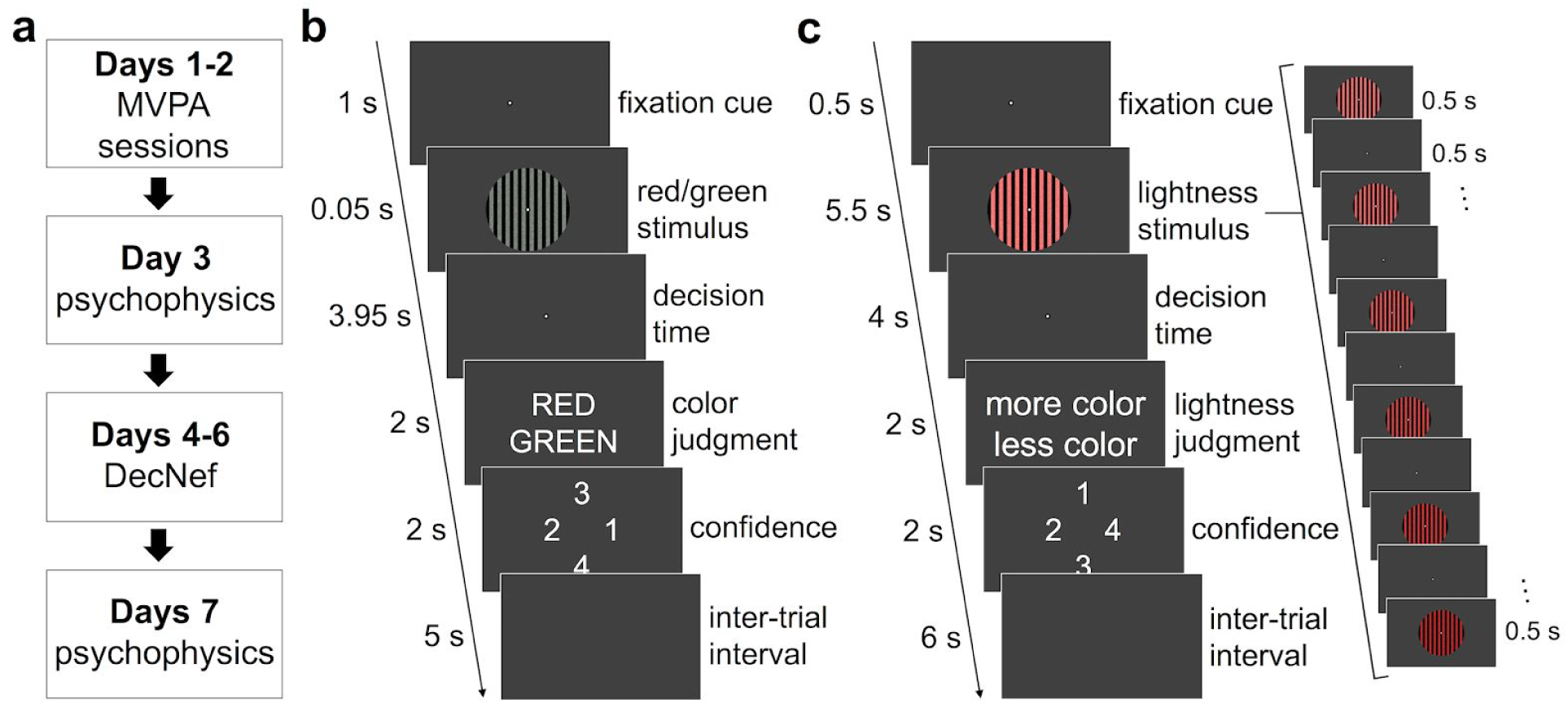
**a)** Experiment flowchart. On Days 1 and 2, participants performed color lightness and color discrimination tasks in the fMRI scanner to decode multivoxel patterns for color and confidence, respectively. On Days 3 and 7 participants performed a red/green color discrimination task outside of the fMRI scanner. On Days 4-6 participants performed a DecNef task in which they were rewarded for the simultaneous activation of multivoxel patterns for red color in visual cortex and high confidence in frontal and parietal cortex. **b)** Confidence decoder task. After a 1 s fixation period, a colored grating (red or green) was presented for 500 ms. After a 1.5 s post-stimulus interval, participants indicated whether they perceived the grating to be red or green and rated confidence on the color discrimination task from 1 (guessed) to 4 (certain). The same task was used in the psychophysics sessions outside of the fMRI scanner on Days 3 and 7. **c)** Color decoder task. Participants viewed 6 circular colored (either all red or all green on a given trial; see Methods) vertical gratings presented for 500ms each and flashed at a frequency of 1 Hz (6 s total). Grating color lightness either increased or decreased (shown) with successive presentations. After a 2 s decision period, subjects indicated whether color lightness increased or decreased by selecting the “less color” or “more color” options, respectively. Text in panels b and c is enlarged compared to its actual size during the experiment for clarity.

### Participants

Seventeen subjects (2 female, mean ± SD age: 26.0 ± 7.5 years, 2 left-handed) with normal or corrected-to-normal vision participated in the decoder construction stage on Days 1 and 2. Two participants were excluded from analyses following the decoder construction stage for not having accuracies greater than 55% for both the color decoder and at least 2 of the 4 confidence decoders (Cortese et al. 2016). Thus, 15 subjects (1 female, mean ± SD age: 25.4 ± 7.1 years, 2 left-handed) are included in the analyses for the behavioral and DecNef tasks on Days 3-7. The study was conducted at the Advanced Telecommunications Research Institute International (ATR) and was approved by the Institutional Review Board of ATR. All subjects gave written informed consent.

### Flicker fusion task

On the first day of the experiment a flicker fusion task (Simonson and Brozek 1952) was used to determine perceptually equiluminant red and green RGB triplets. On each trial of the flicker fusion task, a flickering circle (30 Hz, diameter ~13.5°) alternated between either red and neutral gray (rgb[128 128 128]) (block 1), green and neutral gray (block 2), or red and green (blocks 3 and 4). The screen background in this and all other tasks both inside and outside of the fMRI scanner was a uniform gray (rgb[64 64 64]). Participants were instructed to use button presses in order to minimize the amount of flicker they perceived in the stimulus as follows. On each trial, one of the two colors textures was used as a reference stimulus while the test stimulus, which was always either red or green, had the corresponding red or green channel of its RGB triplet shifted either up or down when participants pressed either the ‘I’ or ‘K’ key, respectively. Participants then pressed the ‘Y’ key to indicate that they had reached a point of minimal flicker. Non-variable RGB channel values in the test texture (e.g., the green and blue channels in the red test texture) were arbitrarily set to 80. On half of the trials the starting value of the variable channel was set to a random value between 0 and 19, while on the other half it was set to a random value between 236 and 255.

There were three practice trials for the flicker fusion task, after which participants completed 4 blocks of 12 trials each. In both the practice and 12-trial blocks, trials were separated by a 2-s intertrial interval (ITI), during which a uniform gray screen was shown. In the first two blocks the test textures were red and green, respectively, and reference textures were neutral gray. In the third block, the reference texture was red with an RGB triplet that corresponded to the mean of all of the 12 selected minimal flicker inducing red RGBs from block 1, while the test texture was green with the same stimulus parameters as the green test textures in block 2. In the fourth block, the reference texture was green with an RGB triplet that corresponded to the mean of all of the 12 selected minimal-flicker inducing green RGBs from block 2, while the test texture was red with the same stimulus parameters as the red test textures in block 1. For each subject, the red and green RGB triplets used throughout the rest of the experiment were computed as the mean of all selected minimal flicker inducing RGBs from blocks 1 and 4 and blocks 2 and 3, respectively (mean ± s.e.m.: red = [218.6 80 80] ± [3.11 0 0], green = [80 149.1 80] ± [0 0.98 0]).

### Red/Green Color Discrimination Task

The red/green color discrimination task (Figure 1b) was performed both outside and inside of the scanner on Days 1 and 2, and outside of the scanner only on Days 3 and 7. At the start of each trial a white fixation circle (diameter ~0.43°) was presented for 1 s on a gray background (rgb[64 64 64]). A circular vertical grating (diameter ~13.5°) and a black annulus (diameter ~0.85°), both centered around the white fixation circle, then appeared for 0.5 s (Figure 1b). The black vertical bars within the grating had a width of ~0.64°, with the area between them subtending the same visual angle.

The majority of pixels in the areas between the black bars had grayscale RGB triplet values (i.e., all RGB channel values were equal) that varied randomly on each frame (frame duration = 16.67 ms) with a mean channel value of 120 and a standard deviation of 51.2. The area between black bars in the grating was thus dynamic. Color strength was adjusted by setting the color of a variable proportion of pixels in the areas between the black bars to either a red or green RGB triplet. Functionally equiluminant RGB triplets determined for each subject in the flicker fusion task were fixed for all red/green color discrimination and color detection tasks used throughout the rest of the experiment. The locations of colored pixels varied randomly between frames, but the proportion of colored pixels was constant throughout a given trial.

Offset of the grating was followed by a 1.5-s decision period in which only the fixation circle remained on the screen. Participants were then asked to report the color of the grating (red or green) and to indicate their confidence in their decision on a scale from 1 to 4 per the following instructions: 1 corresponded to a guess, 2 corresponded to having low but non-zero confidence, 3 corresponded to having moderately high confidence without being certain, and 4 corresponded to feeling certain in their decision. Participants had two seconds to make each response. The on-screen locations of each response option (‘Red’ and ‘Green’ for the color judgment and ‘1’,’2’,’3’, and ‘4’ for the confidence judgment) were randomized on each trial. For all iterations of this task that occurred outside of the fMRI scanner on Days 1 and 2 trial-by-trial feedback (1 s) was given in the form of a green (rgb[0 255 0]) “+1” for correct discrimination responses or a red (rgb[255 0 0]) “-1” for incorrect discrimination responses. The ITIs for this task where 1 s and 5 s when performed outside and inside of the fMRI scanner, respectively.

On both Day 1 and Day 2, prior to the decoder construction session participants performed 80 trials of an adaptive version (QUEST, (Watson and Pelli 1983)) of the red/green color discrimination task. The adaptive procedure used two interleaved 40-trial staircases to estimate the stimulus strength (in proportion of colored pixels) that would lead to 75% correct accuracy on the task. These procedures were broken down into two 40 trial blocks. The mean of the two 75% correct threshold estimates on Day 1 was used as starting stimulus strength for the red/green discrimination task in the subsequent Day 1 decoder construction session in the scanner. The mean of the two 75% correct threshold estimates on Day 2 was used to determine the stimulus strengths that would be used for the pre- and post-DecNef psychophysics tasks on Days 3 and 7 (see below).

### Color Lightness Task

The color lightness task (Figure 1c) was performed both inside and outside of the scanner on Day 1, and inside the scanner on Day 2. On each trial, a fixation circle with the same parameters as that in the red/green color discrimination task appeared for 1 s. A colored grating stimulus (either red or green) then flashed for 0.5 s durations at 1 Hz and its color lightness either increased or decreased linearly over a period of 6 s (6 presentations in total).

The grating stimulus had the same parameters as that in the red/green color discrimination task except for the following. In the area between the black vertical bars, all pixels were colored (either all red or all green). On each trial, a set of 6 different equally spaced values for the dominant RGB channel was determined. This set had a variable range across trials but a constant mean equal to the dominant channel value in the corresponding RGB triplet determined by the flicker fusion task. On each frame of a given 0.5-s grating presentation, the dominant RGB channel value of a given colored pixel was drawn from a normal distribution with a mean of the corresponding set value and a SD of 51.2. The value of the non-dominant RGB channels for a given colored pixel was determined by taking the difference between the dominant and non-dominant rgb channel values from the relevant RGB triplet determined by the flicker fusion task, and subtracting it from the dominant RGB channel value for that pixel. Thus, for each subject the difference between dominant and non-dominant RGB channel values per color was constant across all colored pixels for all grating presentations in this task.

After grating presentation there was a 2-s decision period in which only the fixation circle remained on the screen. Participants then had 2 s to indicate whether the flashing grating stimulus increased or decreased in lightness over time. Because the color tended to look more saturated when lightness decreased, the response options for decreases and increases in lightness were “more color” and “less color”, respectively (Figure 1c).

Participants performed 16 trials of this task outside of the scanner on Day 1. On these lightness trials, but not those performed inside the scanner, they received the same trial-by-trial feedback for correct and incorrect judgments as they did in the color discrimination task. The first four trials were designed to familiarize subjects with the task, and thus each stimulus in these trials used a large range of lightness values (154 RGB units). Trials 5 to 16 employed a 1-up 1-down staircasing procedure with variably weighted step sizes (Kingdom and Prins 2010) of 7.68 and 2.56 RGB units, respectively. The starting range of lightness values on trial 5 was 15 RGB units. The mean of all of the lightness range values from all staircased trials in which a reversal (i.e., a correct response following an incorrect response or vice versa) occurred was set as the midpoint of the uniform distribution of potential range values used in the first color decoder run of the subsequent Day 1 decoder construction session.

### Color and Confidence MVPA Overview

Color and confidence decoders were trained on multivoxel BOLD signal patterns acquired while participants performed the color lightness and red/green color discrimination tasks, respectively. The color decoder was trained on voxel activities in a region of interest (ROI) spanning visual areas V1, V2, V3, and V4 (denoted hereafter as V1-4). Separate confidence decoders were trained on voxel activities in each of four frontoparietal ROIs: inferior parietal lobule (IPL), inferior frontal sulcus (IFS), middle frontal gyrus MFG, and middle frontal sulcus (MFS) (Figure 2). Each task was performed in separate 16-trial runs. Color (lightness task) and confidence (red/green color discrimination task) runs alternated consecutively for each participant, with the order pseudorandomized across participants. Participants performed as many of each run as possible across two 90 minute scanning sessions on Days 1 and 2 (mean ± SD across subjects: 9.5 ± 0.9 color runs and 9.9 ± 0.8 confidence runs).

**Figure 2.**
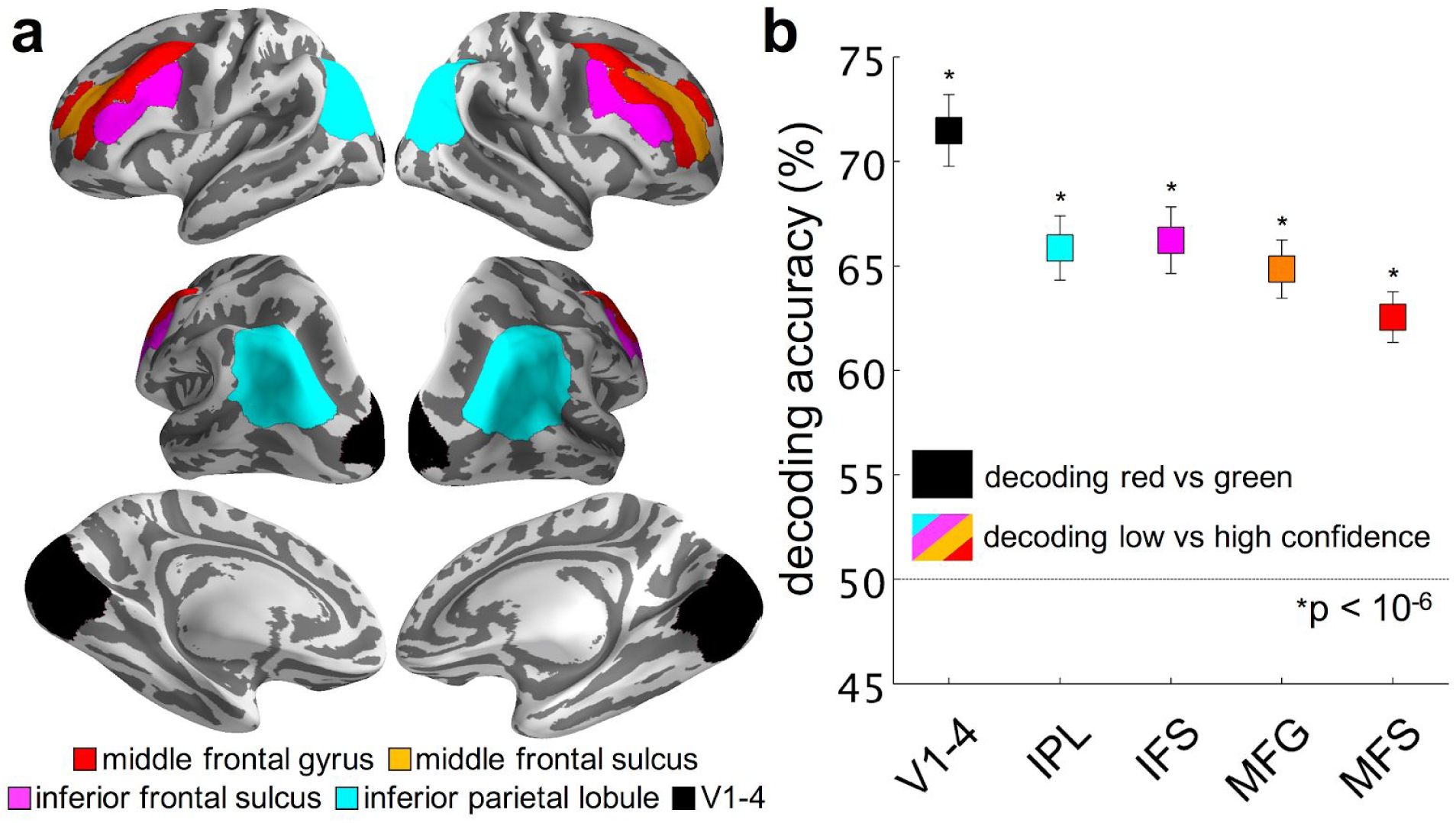
**a)** Average regions of interest (ROIs) used for decoder construction. Voxels were included in each of the displayed average ROIs if they were present for at least half of the 17 decoder construction participants. ROIs are displayed on an average (N=17) inflated cortical surface using the Freesurfer and PySurfer software packages. **b)** Decoding accuracies (reported as mean ± s.e.m., N =17). All decoders were trained using sparse logistic regression (Yamashita et al. 2008) and tested with 10-fold cross-validation. Color decoding (red vs green) accuracy in V1-V4: 71.5 ± 1.7%. Confidence decoding (high vs low) accuracy in IPL: 65.9 ± 1.5%, IFS: 66.2 ± 1.6%, MFG: 64.9 ± 1.4%, MFS: 62.6 ± 1.2%. All decoding accuracies were significantly higher than chance (50% correct) as measured by Bonferroni corrected (α**_corrected_** = 0.01), two-tailed one-sample t-tests. V1-4: combined visual areas V1, V2, V3, & V4; IPL, inferior parietal lobule; IFS, inferior frontal sulcus; MFG, middle frontal gyrus; MFS, middle frontal sulcus.

Iterative sparse logistic regression (Yamashita et al. 2008) was used to select and weight the most informative voxels for distinguishing red vs green color in the visual ROI and high vs low confidence in the four frontoparietal ROIs as previously described (Amano et al. 2016; Cortese et al. 2016). Decoding accuracy was validated using an iterative leave-one-run-out procedure. For each cross validation run, the SLR algorithm selected and weighted a subset of voxels in the relevant ROI. These voxels were then removed, and the algorithm was applied again, selecting and weighting a new, unique subset of voxels. This process was repeated iteratively, 10 times for each cross validation run. Decoding accuracies were then averaged across cross validation runs for each iSLR iteration, and the number of iterations that led to the highest decoding accuracy was selected as the optimal number to be subsequently used during DecNef.

Following the cross validation procedure a separate training run was performed on the entire dataset using the optimal number of iterations. The resulting decoder was used for the subsequent DecNef sessions on Days 4-6. The output of the color decoder reflected the probability of the participant viewing a red stimulus, while the output of the confidence decoder reflected the probability of the participant being in a state of high perceptual confidence.

### MVPA Task Thresholding

For each task performed during the MVPA sessions, in the majority of trials (83.7% ± 0.7% of red/green discrimination trials and 83.6% ± 0.7% of lightness trials), hereafter described as threshold trials, stimulus strength was titrated via a run-by-run thresholding procedure in order to keep performance near 75% correct. This was intended to 1) facilitate a good spread of low to high confidence responses on the red/green discrimination task, and 2) keep participants engaged in the lightness task. The remaining trials either had relatively high stimulus strength (lightness range = 38.4 RGB units in the lightness task, 10.8% ± 0.9% of color trials; 80% of colored pixels for the red/green discrimination task; 10.7% ± 0.7% of confidence trials) or zero stimulus strength (no change in lightness in the lightness task, 5.6% ± 0.1% of color trials; zero colored pixels in the red/green discrimination task, 5.6% ± 0.1% of confidence trials). These high and zero stimulus strength trials were randomly interleaved across runs.

The difficulty of threshold trials in the color decoder task (color lightness) was modulated by changing the range of lightness values across which the colored pixels in the grating stimulus increased or decreased. For a given run, lightness range values were drawn from a uniform distribution, the range of which was arbitrarily set to 3.84 RGB units when the median of the distribution was greater than or equal to 5.12 RGB units, and 150% of the median when the median was less than 5.12 RGB units. The median of this distribution of lightness range values was adjusted per run (with the exception of the first run on Day 1) based on performance in the preceding run according to the following rules. If the percent correct score on threshold trials in the preceding run was greater than or equal to 95%, between 80% and 95%, between 55% and 70%, or less than or equal to 55%, then the median of the current run’s distribution of possible lightness values was scaled by 70%, 80%, 120%, or 130%, respectively.

For the first color decoder run on Day 1, the median of the distribution of possible lightness range values for threshold trials was set to the mean lightness range across all lightness task reversal trials from the Day 1 pre-decoder construction 1-up 1-down lightness task staircasing procedure (see above). The run-by-run thresholding procedure succeeded in maintaining group performance on threshold trials near perceptual threshold (mean ± s.e.m. percent correct = 73.7% ± 1.4%, d’ = 1.53 ± 0.08).

The difficulty of the confidence decoder task (red/green discrimination) was modulated by changing the proportion of colored pixels in the grating stimulus. On the first confidence decoder run of Day 1, the proportion of colored pixels for a given trial was drawn from a uniform distribution, the minimum and maximum of which corresponded to mean Quest-estimated stimulus threshold from the Day 1 pre-decoder construction adaptive staircasing procedure multiplied by 1.2 and 1.6, respectively. The multipliers in this case were both greater than 1 to account for the observation from pilot subjects that the Quest procedure on Day 1 tended to underestimate the color stimulus strength that would lead to 75% correct accuracy on the red/green color discrimination task inside the scanner. On subsequent runs, the range of this distribution was scaled according to the same rules as those in the color decoder task (see above). Group performance on threshold trials in this task was also maintained near perceptual threshold (mean ± s.e.m. percent correct = 74.3% ± 1.57%, d’ = 1.67 ± 0.10, confidence ratings = 2.15 ± 0.14).

### fMRI localizer scans

In order to determine the subregions of V1, V2, V3 and V4 that retinotopically mapped to the grating stimuli in the color and confidence decoder tasks, during the second decoder construction session (Day 2) participants were presented with a flickering colored checkerboard localizer stimulus that occupied the same subregion of the visual field as those grating stimuli (0.425°-6.75° eccentricity). The localizer stimulus was presented alternately, in 8-s periods, with a second flickering colored checkerboard stimulus whose dimensions corresponded to the black annulus inside of the grating stimuli in the color and confidence decoder tasks (0.215°-0.425° eccentricity). Each stimulus was presented 14 times per run (224 s total), and each participant performed 3 runs. To ensure that participants maintained their gaze at the fixation point throughout each run, they performed a change detection task in which they pressed a button every time the fixation point changed color.

### Optimization of Color and Confidence MVPA

Color and confidence decoders were constructed using sparse logistic regression as previously described (Amano et al. 2016; Cortese et al. 2016; Yamashita et al. 2008). To account for variability in hemodynamic delay (Buckner 1998), for each region of interest (ROI), we trained separate decoders for each of several time windows for each type of decoder construction run (color or confidence). All decoding windows were shifted back in time to account for an assumed average hemodynamic delay of 6 seconds. In what follows, we indicate the time window according to the event to which the time window is supposed to correspond assuming this 6-s shift; e.g., when we indicate that a time window started at the target stimulus onset, that means that the first fMRI image that was analyzed in that time window was the one that was captured 6 seconds after induction cue onset.

For color runs, where the target stimulus was flashed over a period of 6 s, we trained decoders over nine different time windows: four 2-s windows starting at timepoints −2 s, 0 s, +2 s, and +4 s relative to target stimulus onset, three 6-s windows starting −2 s, 0 s, and +2 s relative to target stimulus onset, and two 8-s windows starting at −2 s and 0 s relative to target stimulus onset, where negative and positive numbers correspond to earlier and later time points, respectively. For confidence runs, which had a much briefer target stimulus display time (0.5 s) we trained decoders over three different time windows: two 2-s windows starting at timepoints −2 s and 0 s relative to target stimulus onset, and one 3-s windows starting −2 s relative to target stimulus onset. Further, for color runs, two decoders were trained over each time window, one in which the V1-4 ROI was intersected with the functional localizer ROI, and one in which it was not. This led to a total of 18 different localizer/time window combinations for color decoding.

For each decoder type for each participant, the time window-localizer combination for color decoding and time window for confidence decoding that resulted in the highest cross-validated accuracy was used for training the corresponding DecNef decoder on the entire dataset (summarized in Table 1). In all cases, the BOLD signal was averaged across all time points within a given time window, and the resulting averaged samples were used for the iSLR decoder training procedure.

**Table 1.** Subject-specific temporal windows and V1-4 localizer intersection status that led to maximum decoding accuracy. A Y in the second column indicates that the maximum decoding accuracy was obtained when the V1-4 ROI was intersected with the functional localizer ROI, while an N indicates that the maximum decoding accuracy was obtained when the entire V1-4 ROI was used. Negative and positive numbers in columns 3 and 5 indicate temporal window starting times before and after target stimulus onset, respectively. The decoding parameters shown here were used to train the decoders that were subsequently used for neurofeedback.

| Participant No. | Localizer | Color Decoder Time Window |  | Confidence Decoder Time Window |  |
| --- | --- | --- | --- | --- | --- |
| | | Start time relative to target stimulus onset (Δs) | Duration (s) | Start time relative to target stimulus onset (Δs) | Duration (s) |
| 1 | N | -2 | 6 | -2 | 6 |
| 2 | N | 0 | 8 | 0 | 4 |
| 3 | N | 0 | 8 | 0 | 4 |
| 4 | N | 0 | 8 | -2 | 6 |
| 5 | Y | 0 | 8 | -2 | 6 |
| 6 | N | 2 | 6 | 0 | 4 |
| 7 | N | 0 | 8 | 0 | 4 |
| 8 | N | 0 | 8 | 0 | 4 |
| 9 | Y | 0 | 6 | 0 | 4 |
| 10 | N | 0 | 8 | 0 | 4 |
| 11 | N | 2 | 4 | 0 | 4 |
| 12 | N | 2 | 4 | 0 | 4 |
| 13 | N | -2 | 8 | 0 | 4 |
| 14 | Y | 4 | 4 | -2 | 6 |
| 15 | N | -2 | 8 | 0 | 4 |
| 16 | N | 2 | 4 | -2 | 4 |
| 17 | Y | 0 | 6 | 0 | 4 |

For confidence decoding, to ensure an equal number of samples in the low and high confidence classes for decoder construction we used the following downsampling approach. Confidence ratings of 1 and 4 were always allocated to the low and high confidence training classes, respectively. The classes to which confidence ratings of 2 and 3 were allocated were determined so as to minimize the difference in the sample number between the two classes. Thus, confidence ratings could be divided into low and high confidence classes in the following three ways: low confidence = ratings of 1, high confidence = ratings of 2-4 (N = 9), low confidence = ratings of 1 and 2, high confidence = ratings of 3 and 4 (N = 6), and low confidence = ratings of 1-3, high confidence = ratings of 4 (N = 2). After assigning confidence ratings to their respective decoder classes, the class with the higher number of training samples was downsampled to equate the total number of samples between classes. The to-be-removed samples were chosen randomly, over four separate iterations. Cross-validated decoding accuracies were calculated for each downsampling iteration as described above, and the training samples that were used in the iteration that resulted in the highest decoding accuracy were subsequently used for training the DecNef decoder.

### DecNef sessions

All participants in the MVPA session who had accuracies of 55% or higher for color decoding and for at least two of the frontoparietal ROIs for confidence decoding (N=15) were included in the DecNef sessions on Days 4-6. Each DecNef run (mean ± s.e.m. = 9.6 ± 0.4 runs per day) started with an initial 29 second fixation period, during which a white fixation cross (diameter ~0.84°) was presented at the center of the screen. This was followed by 16 trials in which participants were rewarded for activating the patterns identified in the MVPA session as corresponding to red in V1-V4 and high confidence in the four frontoparietal ROIs (Figure 3a). On each trial, after a 1 s cue, participants viewed a vertical grating with the same dimensions as the gratings shown during the MVPA session for 6 s, during which time they were instructed to try to use their minds to activate a pattern of brain activity in order to make the size of a subsequent feedback stimulus (a black disc) as large as possible. The feedback disc appeared for 2 s after a 6 s rest period (Figure 3a).

**Figure 3.**
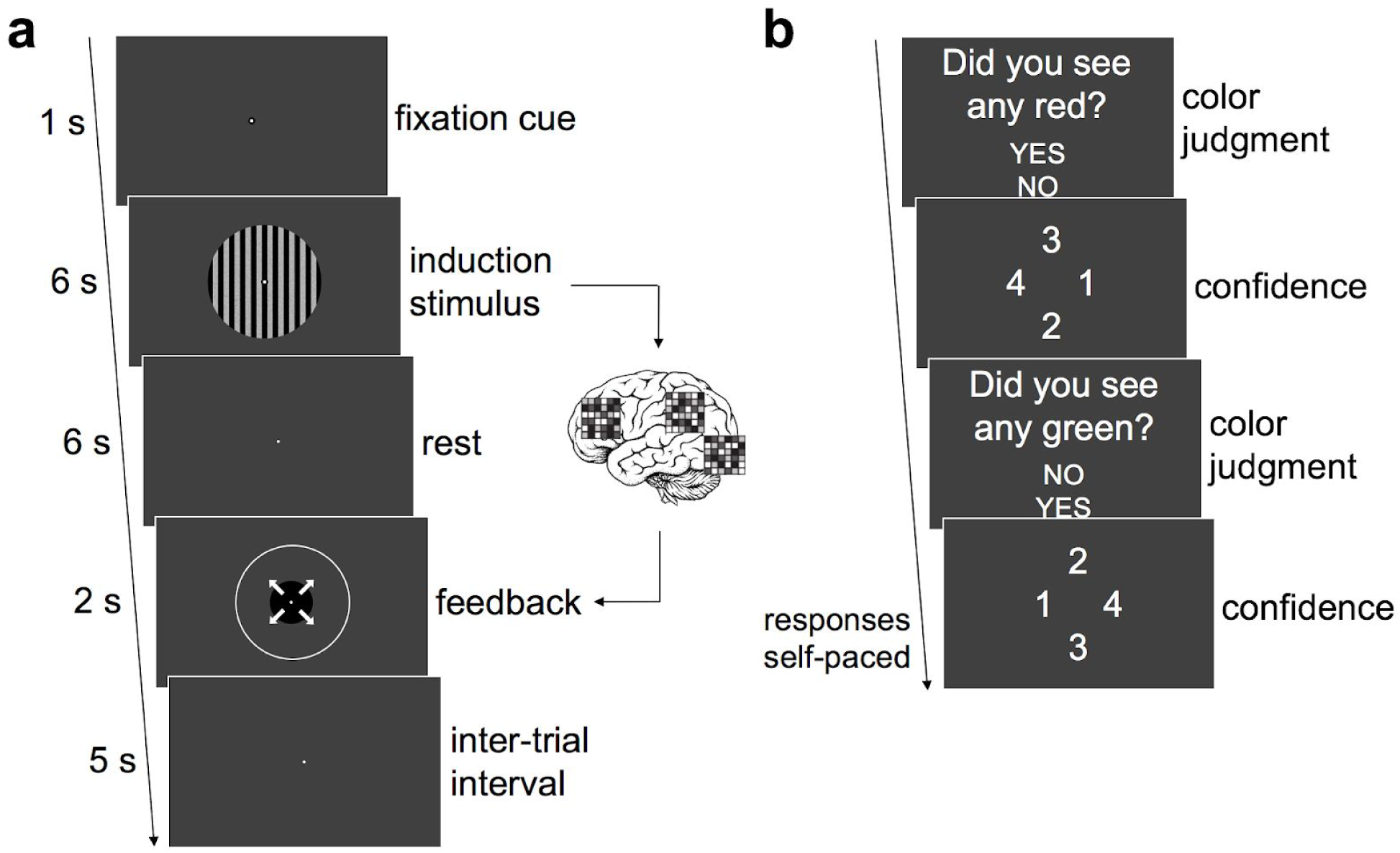
DecNef task. **a)** Trial structure. Participants were told that, after a 1 s cue, while an induction stimulus (vertical grating) was present, they should try to activate a pattern of brain activity so as to maximize the size of a subsequently presented black feedback disc. A 6 s rest period separated the induction and feedback stimuli to account for hemodynamic delay and real-time processing of fMRI images. BOLD signal in the visual (V1-4) and frontoparietal (IPL, IFS, MFG, MFS) ROIs from the induction period was processed by the previously-trained color and confidence decoders, respectively. The magnitude of the resulting red and high confidence likelihoods determined the size of the feedback disc such that participants were maximally rewarded for simultaneously activating a red pattern in visual cortex and a high confidence pattern in frontoparietal cortex. **b)** End of run questions. At the end of each run participants were asked whether they perceived red or green in the induction grating during any of the 16 trials in that run. They were also asked to rate confidence in each of these judgments on a scale from 1 (low) to 4 (high). Text in panel b is enlarged compared to its actual size during the experiment for clarity. V1-4: combined visual areas V1, V2, V3, & V4; IPL, inferior parietal lobule; IFS, inferior frontal sulcus; MFG, middle frontal gyrus; MFS, middle frontal sulcus.

For online decoding, the BOLD signal was head motion corrected in real time using Turbo-Brian Voyager software (Brain Innovation, Netherlands). The BOLD signal corresponding to the interval from the start of the run until the last measured TR in the prevailing trial was then extracted from the voxels that were selected in each ROI during the MVPA session. Linear detrending and z-score normalization was then performed on these extracted voxel activities. The resulting detrended, z-score normalized signal in each ROI was then averaged across the 6 second rest period of the prevailing trial, which should correspond to neural activity during the 6 second induction period when adjusting for an estimated 6-s hemodynamic delay, and was inputted into the corresponding color or confidence decoder. The resulting decoding likelihoods (LLs) determined the size of the feedback disc according to the following formula:

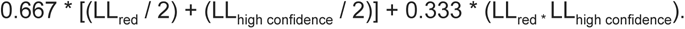

The size of the feedback disc also corresponded to a monetary reward earned on each trial (max = 18.75 yen or approximately $0.15 US dollars per trial). Successive trials were separated by a 5 s ITI.

To ensure that the correct voxels were targeted during DecNef, we computed the correlation between the detrended, z-score normalized signal in each ROI during DecNef and the mean detrended, z-score normalized signal in the corresponding ROIs across all decoder construction trials for each subject. Any trial that resulted in a correlation value less than r = 0.6 in any of the five target ROIs was excluded from the current analyses [median (interquartile range) = 1.3% (0.5% − 4.3%) of trials excluded per participant].

For all but two DecNef participants, a small proportion of DecNef trials [median (interquartile range) = 1.1% (0.4% − 1.8%) of total DecNef trials per participant (N=13)] motion-corrected BOLD signal data could not be retrieved in time for feedback stimulus presentation. On these trials a blank gray screen was presented during the feedback interval (Figure 3a). Participants were instructed beforehand that any such trials would be the result of computer malfunctions and should be ignored. These trials were omitted from all data analyses.

On achromatic trials, normally distributed gray RGB triplets for pixels between the black vertical bars in the induction stimulus were generated as described for the red/green color discrimination task above. To generate red and green induction stimuli, the same procedure was followed, but a value of 12.8 was either subtracted from the green and blue channels or added to the green channel, respectively, of each dynamic voxel.

At the end of each DecNef run participants separately reported whether they perceived any red or any green in the induction grating stimulus on any of the 16 trials in that run, and indicated how confident they were in this judgment on the same 1 to 4 scale that was used during decoder construction trials (Figure 3b). The order in which the red and green perception questions were asked at the end of each DecNef run was randomized across runs. Importantly, on 97.4 ± 0.2% of trials, the induction stimulus was achromatic, while on the remaining trials (4 per day, the induction stimulus was either slightly red (2 trials) or slightly green (2 trials). Specifically, on each day of neurofeedback one run contained one red trial, a different run contained one green trial, and a third run contained both one red and one green trial. Run order was randomized between subjects, but the three runs containing color trials were constrained to always occur within the first 8 runs on a given day to avoid a given subject missing a run with a color trial due to time constraints.

Given this setup, each run can be categorized as one of four classic types according to signal detection theory: hits (reported seeing color during a run in which at least one trial contained a colored induction stimulus), misses (reported seeing no color during a run in which at least one trial contained a colored induction stimulus), false alarms (reported seeing color during a run in which no trials contained a colored induction stimulus), and correct rejections (reported seeing no color during a run in which no trials contained a colored induction stimulus). The color manipulation was designed to induce a nonzero baseline false alarm rate for perceiving color, e.g., reporting the perception of red in a given DecNef run when no red was present in any induction stimuli during that run.

At the end of each DecNef session participants were asked what strategies, if any, they used during neurofeedback to try to maximize the size of the feedback circle. At the end of Day 7, participants were asked two debriefing questions. First, they were asked whether they thought they received real or sham neurofeedback. Second, they were asked to guess, assuming they had been receiving real neurofeedback, whether they were rewarded for activating a pattern of brain activity corresponding to red perception or green perception.

### Pre-/post-DecNef color discrimination task

On Days 3 and 7 participants (N=15) performed the same red/green color discrimination task from Days 1 and 2, with the following differences in stimulus parameters. Three stimulus levels (proportion of colored pixels), fixed across color to preserve equiluminance, were used to target percent correct scores of 65%, 75%, and 85%. As in the red/green color discrimination tasks performed outside of the scanner on Days 1 and 2, the ITI was 1 s.

Participants first performed 10 practice trials with trial-by-trial feedback (as described in the Red/Green Color Discrimination Task section above). They then performed 6 blocks of 51 trials each with self-paced breaks between blocks and no trial-by-trial feedback. Of the 306 total trials, 276 had stimulus strengths near perceptual threshold, with 46 trials at each of the three near-threshold stimulus strengths for each color. Of the remaining 30 trials, 15 had a high percentage of colored pixels (45%), which was intended to help maintain perceptual templates for color, and 15 had zero colored pixels. All trial types were randomly interleaved across blocks.

The three stimulus strengths were determined for each participant by multiplying their mean Quest-estimated threshold stimulus strength (in proportion of colored pixels) from Day 2 by three proportions (mean ± SD proportions across subjects = 0.57 ± 0.17, 1.06 ± 0.10, 1.56 ± 0.16 for low, medium, and high stimulus strengths, respectively). These proportions were adjusted on a subject-by-subject basis according to the Quest procedure’s tendency to over- or underestimate threshold stimulus strength when considering all of the across-subject data that had been collected at the time. The resulting mean ± s.e.m. performance scores across Days 3 and 7 for low, medium, and high stimulus strengths were 65.0 ± 2.2%, 77.0% ± 2.7%, and 84.0% ± 2.4% correct (d’ = 1.00 ± 0.16, 1.83 ± 0.23, and 2.37 ± 0.22, respectively).

Individual participant data from each day were fit with cumulative normal psychometric functions with free parameters α (threshold) and β (slope), and fixed parameters γ (lapse rate) = 0 and δ (guess rate) = 0 also using the Palamedes toolbox (Prins and Kingdom 2018; Kingdom and Prins 2010). Values on the abscissa were equated across subjects to equal ±1, ±2, and ±3 to reflect the low, medium, and high stimulus strength conditions for each color, respectively, with positive and negative values corresponding to red and green stimuli, respectively. The point of subjective equality (PSE), which corresponds to the stimulus strength at which participants are equally likely to choose red or green on the color discrimination task (i.e., 50% on the ordinate), was estimated as the threshold parameter, α, from the fitting procedure. Given that stimulus strength values were equated across subjects for psychometric curve fitting, the resulting mean ± SD PSE values (PSE_pre-DecNef_ = 1.66 ± 2.00, PSE_post-DecNef_ = 0.34 ± 0.72) can be thought of as the proportion of the lowest stimulus strength necessary for the stimulus to be equally likely to have a subjective appearance of redness or greenness, with positive values reflecting red stimulus strength and negative values reflecting green stimulus strength.

### Apparatus

Stimuli for tasks performed outside of the fMRI scanner were presented on an IBM P275 CRT monitor with a 1280 x 960 resolution and a 60 Hz refresh rate. All visual stimuli were generated with custom Matlab R2014a (Natuck, MA) scripts using PsychToolbox 3.0.12. Stimuli for tasks performed inside of the fMRI scanner were presented on an LCD projector that also had a 1280 x 960 resolution and a 60 Hz refresh rate. Repeated measures ANOVAs were performed using SPSS v22 and were adjusted for violations of the assumption of sphericity with the Greenhouse-Geisser correction when necessary.

### MRI Parameters

MRI images were acquired using 3T MRI scanners (Siemens, Verio [N=15] or Siemens, Trio [N=2]) at the ATR Brain Activation Imaging Center. Both scanners used head coils. Functional images for MVPA and DecNef sessions were acquired using gradient EPI sequences with 33 contiguous slices (repetition time (TR) = 2 s, echo time (TE) = 26 ms, flip angle = 80 deg, voxel size = 3 x 3 x 3.5 mm^3^, 0 mm slice gap) oriented parallel to the AC-PC plane, covering the entire brain. T1-weighted MR images (MP-RAGE; 256 slices, TR = 2s, TE = 26 ms, flip angle = 80 deg, voxel size = 1 x 1 x 1 mm^3^, 0 mm slice gap) were also acquired during the first MVPA session. These images were used for automatic brain parcellation in Freesurfer (Fischl et al. 2002).

### fMRI preprocessing

fMRI images from decoder construction sessions were preprocessed as previously described (Cortese et al. 2016). T1-weighted structural images were processed with an automatic parcellation procedure based on volumetric segmentation and cortical reconstruction using the FreeSurfer image analysis suite (http://surfer.nmr.mgh.harvard.edu/). The IPL, IFS, MFG, MFS ROIs (Figure 2) used in subsequent analyses were defined using this procedure. Visual ROIs were defined using a probabilistic atlas (Wang et al. 2015). Average inflated cortical surfaces shown in Figure 2 were generated using Freesurfer and displayed using PySurfer (https://pysurfer.github.io/). Average ROIs in Figure 2 were generated in Freesurfer for display purposes; voxels were included in each average ROI if they were present in the individual ROIs of at least half of the 17 decoder construction participants (Figure 2a). Gray matter masks were generated using the mrVista software package for Matlab (http://vistalab.stanford.edu/software/), which uses functions from the SPM suite (http://www.fil.ion.ucl.ac.uk/spm), to ensure that only gray matter voxels were used for subsequent analyses. Three-dimensional rigid-body motion correction was applied in mrVista to align functional scans to the T1-weighted structural image for each participant. Day 2 Localizer scans were slice-time corrected and averaged across stimulus groups, and a coherence analysis was applied to identify voxels in visual cortex that responded maximally to the localizer stimulus (Wandell and Winawer 2011). No temporal or spatial smoothing was applied. For all color and confidence decoder construction scans, we removed voxels with exceptional values, extracted BOLD signal time courses from each remaining voxel in each ROI, applied linear detrending, and z-score normalized the BOLD signal per run to account for potential baseline differences between runs.

## Results

### Decoding color and confidence

Based on 10-fold cross validation, color decoding accuracy (mean ± s.e.m.) in visual cortex was 71.5 ± 1.7%, while the mean confidence decoding accuracy across frontoparietal ROIs was 64.9 ± 1.4% (IPL: 65.9 ± 1.5%, IFS: 66.2 ± 1.6%, MFG: 64.9 ± 1.4%, MFS: 62.6 ± 1.2%; Figure 2b). Decoding accuracy in each ROI was significantly greater than chance (50% correct) [V1-4: t(16) = 12.5, p < 0.001, 95% CI = (0.68,0.75); IPL: t(16) = 10.3, p < 0.001, 95% CI = (0.63,0.69); IFS: t(16) = 10.2, p < 0.001, 95% CI = (0.63,0.70); MFG: t(16) = 10.6, p < 0.001, 95% CI = (0.62,0.68); MFS: t(16) = 10.3, p < 0.001, 95% CI = (0.60,0.65); Bonferroni corrected (α**_corrected_** = 0.01), two-tailed one-sample t-tests]. The mean ± s.e.m. numbers of selected voxels in each ROI were the following: V1-V4 = 140.4 ± 23.0, IPL = 107.4 ± 13.1, IFS = 71.5 ± 12.1, MFG = 80.4 ± 14.6, MFS = 79.1 ± 12.6). Two participants were excluded from subsequent DecNef analyses for failing to have accuracies greater than 55% for color decoding and for at least two of the four frontoparietal confidence decoders.

### Decoded Neurofeedback

The color manipulation in the induction stimulus succeeded in establishing a non zero baseline false alarm rate (FAR) for red perception in 9 of 15 DecNef participants (FAR_red_ = 14.2% ± 2.2%) and for green perception in 14 of 15 DecNef participants (FAR_green_ = 23.0% ± 5.9%). One participant was excluded from analyses of false alarm and correct rejection runs because they did not make any false alarms. One additional participant was excluded from these analyses due to a failure to make correct rejections for *both* red and green responses on any single run. Thus, in 13 DecNef participants, we could analyze whether there was any connection between activation of multivoxel patterns for color or confidence and false color perception by comparing color and confidence induction likelihoods between false alarm and correct rejection runs.

In subjects who made red false alarms, there was no significant difference in color induction likelihoods between red false alarm runs and correct rejection runs [t(8) = −0.34, p = 0.75, 95% CI = (−0.24, 0.18), two-tailed paired-samples t-test; Figure 4a]. In subjects who made green false alarms there was no significant difference in color induction likelihoods between green false alarm runs and correct rejection runs [t(12) = −0.34, p = 0.74, 95% CI = (−0.08, 0.06), two-tailed paired-samples t-test; Figure 4a]. However, collapsing across color, high confidence induction likelihoods were significantly higher during false alarm runs than they were during correct rejection runs [t(12) = 2.75, p = 0.02, 95% CI = (0.01,0.06), two-tailed paired-samples t-test; Figure 4b].

**Figure 4.**
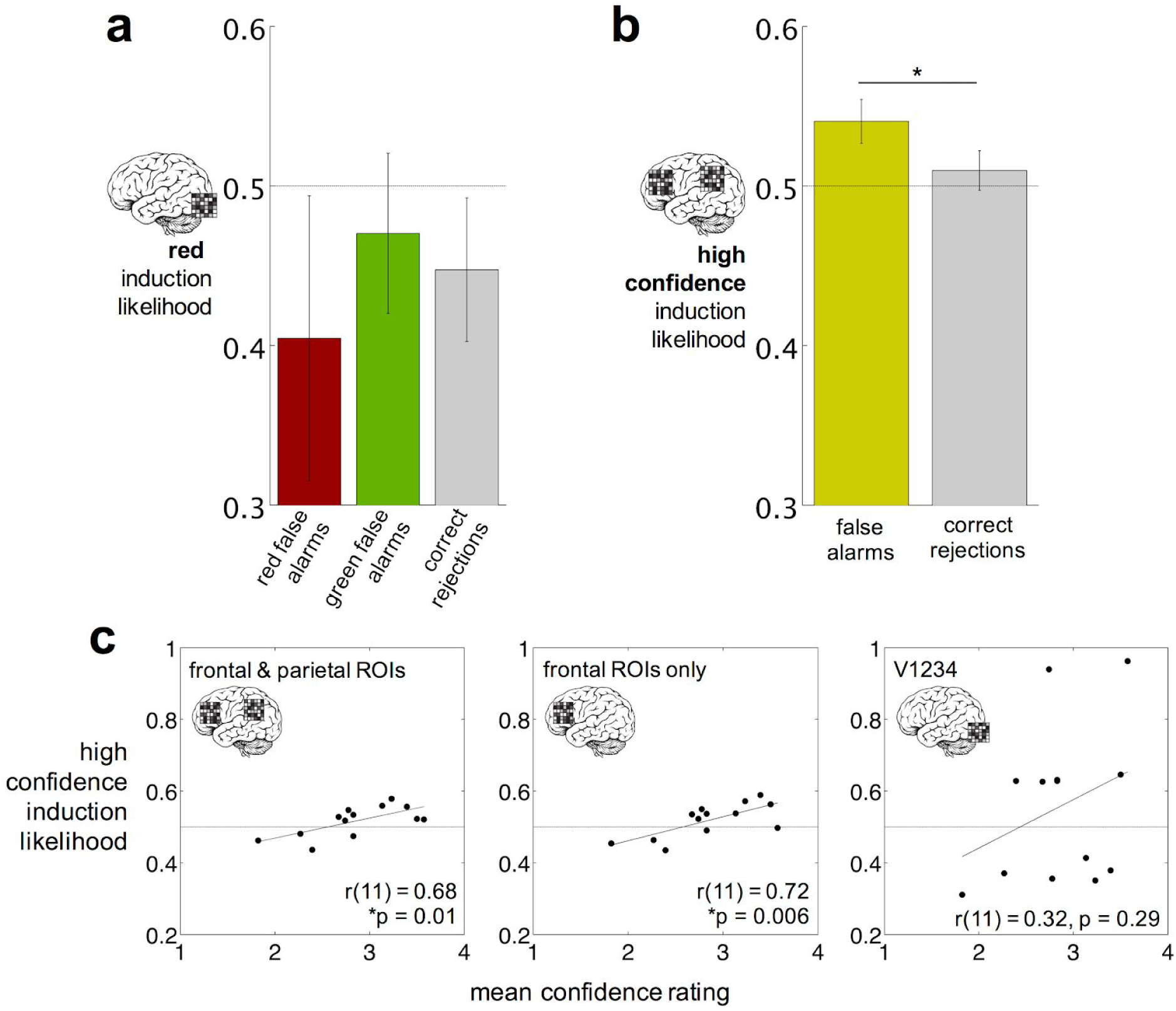
Induction likelihoods during false alarm vs correct rejection runs and pre-/post-DecNef psychometric functions for color discrimination. **a)** Red induction likelihoods during red false alarm, green false alarm, and correct rejection runs. Induction likelihoods were not significantly different between false alarms and correct rejections for either color [red false alarm versus correct rejection runs: t(8) = −0.34, p = 0.75, 95% CI = (−0.24, 0.18); green false alarm versus correct rejection runs: t(12) = −0.34, p = 0.74, 95% CI = (−0.08, 0.06), two-tailed paired-samples t-tests]. **b)** High confidence induction likelihoods during false alarm runs (collapsed across color) and correct rejection runs. High confidence induction was higher under false alarm runs than it was under correct rejection runs [t(12) = 2.75, p = 0.02, CI = (0.01,0.06)]. **c)** Relationship between mean high confidence induction likelihoods and mean decnef confidence ratings across runs. High confidence induction likelihoods and confidence ratings were averaged across all DecNef runs for each subject. Confidence ratings were averaged across color for each DecNef run. The confidence decoder in V1234 was trained in the same manner as those in the frontoparietal ROIs, but was not used for neurofeedback. Bonferroni corrected Pearson correlations suggest a relationship between DecNef confidence ratings and high confidence induction likelihoods in the collective frontoparietal ROI (left panel: IPL + IFS + MFS + MFG, r(11) = 0.68, p = 0.01), and when looking at only the frontal ROIs alone (middle panel: IFS + MFS + MFG, r(11) = 0.72, p = 0.006), but not in visual cortex (right panel: V1-4, r(11) = 0.32, p = 0.29). IPL, inferior parietal lobule; IFS, inferior frontal sulcus; MFG, middle frontal gyrus; MFS, middle frontal sulcus.

If the learned co-activation of the decoded red and high confidence patterns can influence conscious visual perception in real time, then we might predict the false alarm rate (FAR) for red perception during neurofeedback to increase across the three days of neurofeedback. However, a repeated measures ANOVA with within-subjects factor time (3 days) showed that the red FAR did not increase over time [F(1.23,17.21) = 0.55, p = 0.50]. Correspondingly, a repeated measures ANOVA with a within-subjects factor of time suggested that neither red nor high confidence induction likelihoods increased over the three days of DecNef (red induction: F_2,28_ = 0.63, p = 0.54; high confidence induction: F(2,28) = 0.79, p = 0.46). Mean induction likelihoods across days were 0.47 ± 0.04 and 0.52 ± 0.01 for red and high confidence induction, respectively. Because target induction likelihoods did not show a significant increase over time, we are precluded from addressing whether this specific type of DecNef learning can affect real-time color perception. However, this does not preclude reinforcement learning based on spontaneous pairings of the target induction pattern, the induction stimulus, and reward (Amano et al. 2016; Shibata et al. 2019; see also Discussion).

It was also found that high confidence induction likelihoods in confidence ROIs averaged across all neurofeedback runs were positively correlated with average confidence judgments (Figure 4c). To see whether this result extended to visual cortex, an additional confidence decoder was trained using the visual ROI (V1-V4). Confidence decoding accuracy in this ROI was significantly greater than chance by a one-sample, two-tailed t-test [mean s.e.m. decoding accuracy = 65.5 ± 1.8%; t(16) = 8.61, p < 0.001, CI = (61.7, 69.3)]. However, induction of high confidence patterns in this ROI did not correlate with actual confidence judgments during neurofeedback as was the case with the frontoparietal ROIs [r(11) = 0.32, p = 0.29; Figure 4c, right].

Examining the contribution of the individual frontoparietal ROIs to the observed association between high confidence induction and false color perception revealed that the effect was driven by activity in the prefrontal ROIs (Figure 5a). To further investigate whether the information in the ROI-specific decoders was shared with other ROIs during neurofeedback, we performed an information leak analysis as previously described (Shibata et al. 2011; Cortese et al. 2016); Figure 5b). Briefly, this analysis quantifies the extent to which multivariate BOLD patterns in one ROI can predict the output of a decoder trained in another ROI. Figure 5b shows minimal information leak between frontal and visual ROIs, with more intermediate information leak between parietal and frontal and parietal and visual ROIs.

**Figure 5.**
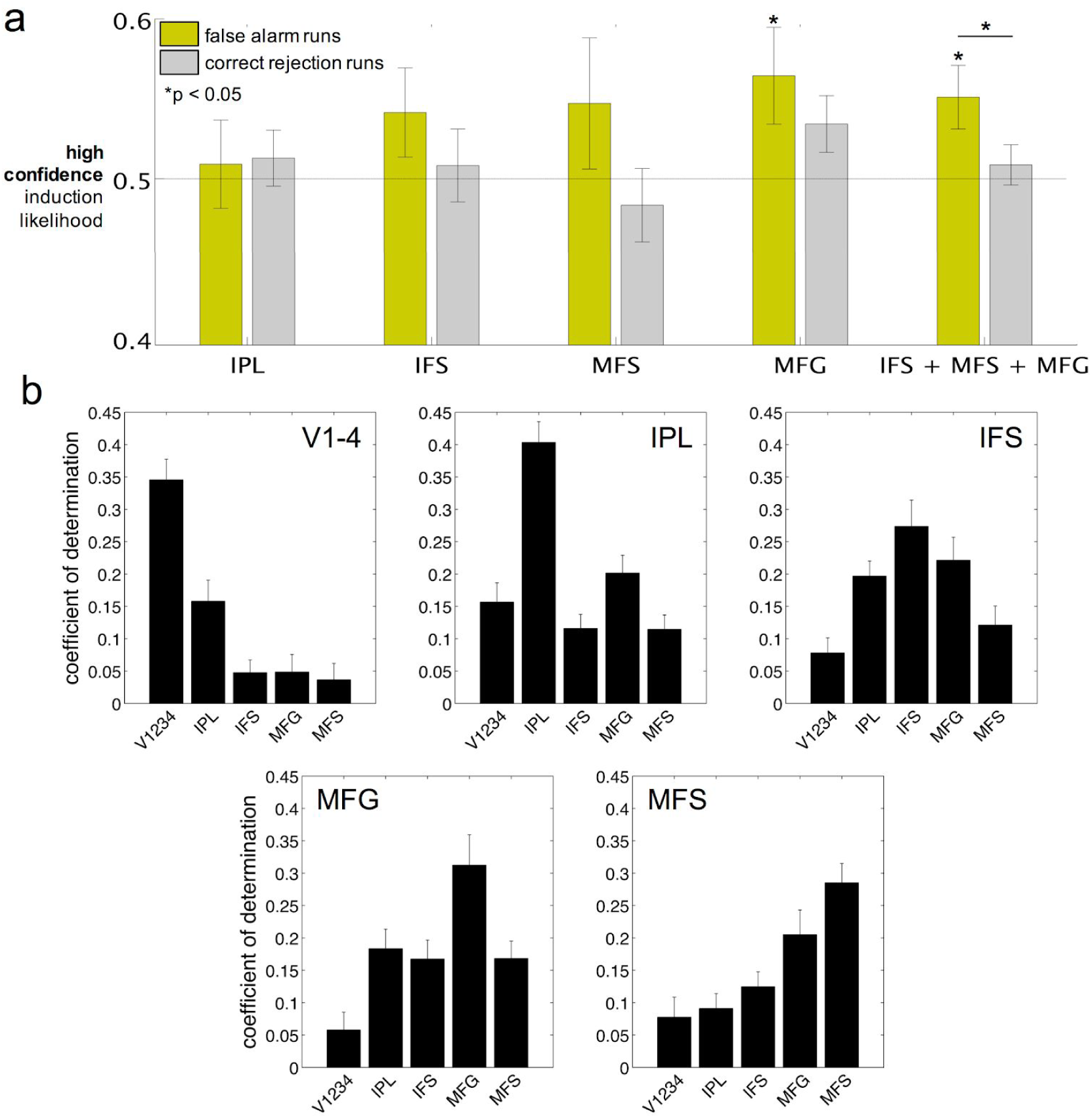
Contributions of individual ROIs. **a)** High confidence induction likelihoods during false alarm vs correct rejection runs in individual prefrontal and parietal ROIs and one group prefrontal ROI (N=13). While the difference in induction likelihoods between false alarm and correct rejection runs in the IFS + MFS + MFG ROI does not survive Bonferroni correction (family-wise alpha for comparing high confidence likelihoods between false alarm and correct rejection runs in each ROI = 0.01), the pattern of results suggests that the difference in high confidence induction likelihoods between false alarm and correct rejection runs found across all frontoparietal ROIs (Figure 4b) is primarily driven by the prefrontal ROIs. *p < 0.05. **b)** Information leak analysis (N=13). The coefficient of determination (y-axis) is an index of the extent to which voxel activities in a given “predictor” ROI (x-axis) can predict, via sparse linear regression (SLiR), color induction likelihoods in V1-4 and confidence induction likelihoods in IPL, IFS, MFG, & MFS. The results show minimal “leak” of information outside of target regions, suggesting that induction likelihoods in a given ROI were minimally influenced by the activities of voxels in neighboring ROIs. This relationship is particularly pronounced when looking at “leak” between ROIs in frontal and visual cortices. V1-4: combined visual areas V1, V2, V3, & V4; IPL, inferior parietal lobule; IFS, inferior frontal sulcus; MFG, middle frontal gyrus; MFS, middle frontal sulcus.

### Pre-/post-DecNef color discrimination

All DecNef participants (N=15) also performed a red/green color discrimination task outside of the fMRI scanner on the day before the first DecNef session and on the day after the last DecNef session. The color discrimination task was the same as that used for confidence decoding in the MVPA sessions (Figure 1c) except that three constant stimulus strengths were used for each color (whereas stimulus strength was modified on a per run basis during the MVPA sessions; see Methods). The resulting individual participant data from each day were fit with cumulative normal psychometric functions (Figure 6). In line with a previous DecNef study of color perception (Amano et al. 2016), participants were significantly more biased towards choosing red after DecNef, as indicated by a significant leftward shift in the point of subjective equality (PSE) [t(14) = 2.43, p = 0.03, 95% CI = (0.16, 2.48), two-tailed paired-samples t-test; Figure 6].

**Figure 6.**
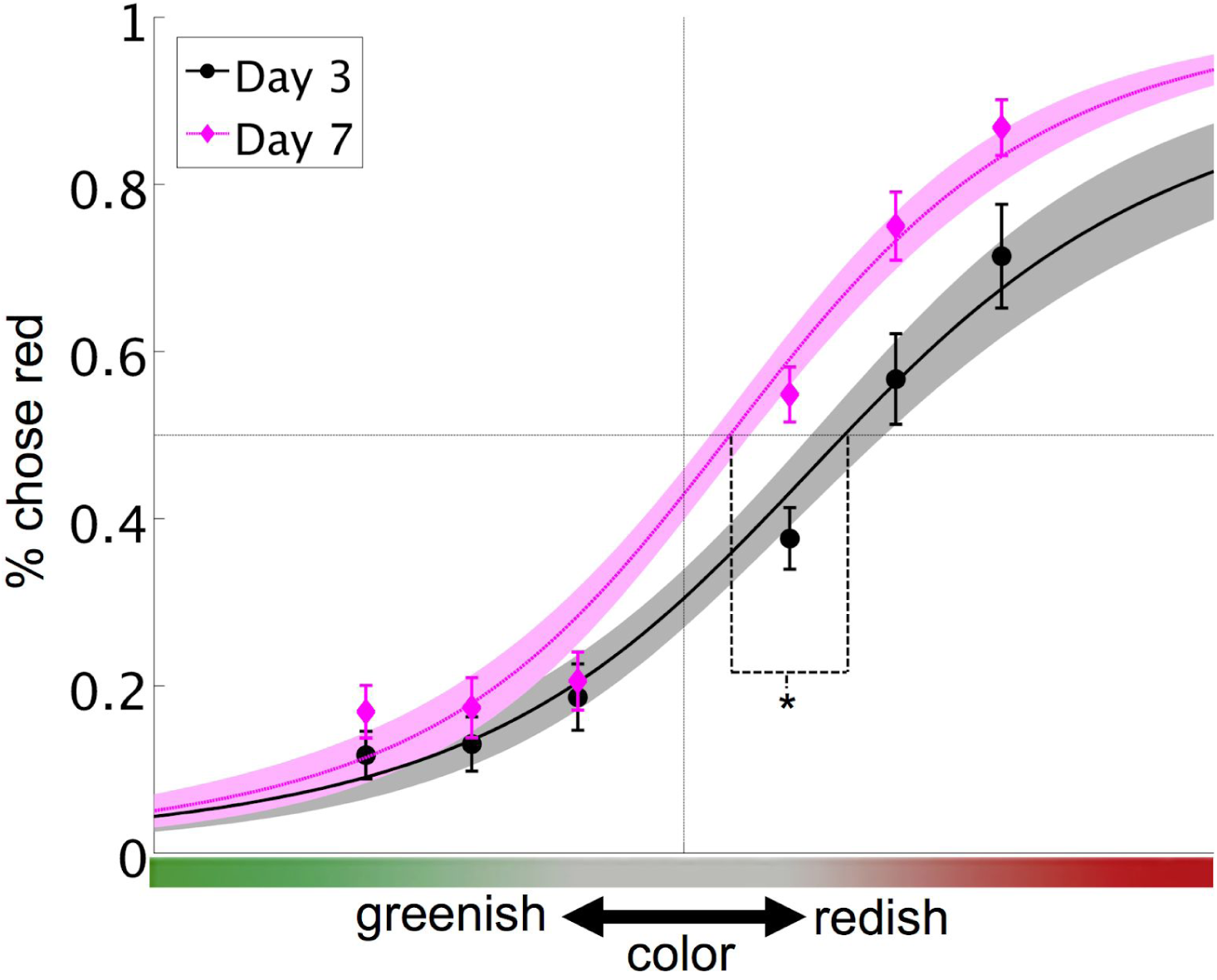
Pre- vs post-DecNef psychometric curves. Participants performed a red/green color task (see Fig. 3b) with the method of constant stimuli on the day before the first DecNef session (Day 3, gray) and on the day after the last DecNef session (Day 7, purple). Psychometric curves were fit to individual participant data using cumulative normal distribution functions. Shown is the mean (black/magenta line) ± s.e.m. (gray/light magenta shaded area) of the individual fits. A significant negative shift in group mean point of subjective equality (PSE) was observed from Day 3 to Day 7 [t(14) = 2.43, *p = 0.03, two-tailed paired-sample t-test], showing a post-DecNef reduction in an initial bias toward choosing green.

The signal detection theoretic measures d’ and criterion (Green DM 1966; MacMillan and Creelman 2004), c, were computed for the color discrimination task with Hits corresponding to trials in which participants correctly discriminated red stimuli as red and False Alarms corresponding to trials in which participants incorrectly discriminated green stimuli as red. There was a trend, though non-significant, toward an overall improvement in discrimination d’ from Day 3 to Day 7 [d’_pre_ = 1.48 ± 0.18, d’_post_ = 1.79 ± 0.22, t(14) = −1.91, p = 0.08, 95% CI = (−0.66, 0.04), two-tailed paired-samples t-test] which may be attributable to perceptual learning (Dosher and Lu 2017). There was also a non-significant trend towards a decrease in criterion [c_pre_ = 0.50 ± 0.12, c_post_ = 0.20 ± 0.08, t(14) = 1.99, p = 0.07, 95% CI = (−0.02, 0.61), two-tailed paired-samples t-test], which is consistent with the observed negative shift in PSE from the psychometric function analyses. There was no significant change in red/green color discrimination confidence from Day 3 to Day 7 [mean confidence_pre_ = 2.13 ± 0.14, mean confidence_post_ = 2.18 ± 0.13, t(14) = −0.54, p = 0.60, 95% CI = (−0.28,0.17), two-tailed paired-samples t-test].

### Controlling for color-confidence biases

Because the confidence decoder was trained on confidence responses from a red/green discrimination task, one concern is that the confidence decoder may be confounded with color. Figure 7a shows that indeed confidence responses were higher on average for green stimuli compared to red stimuli in all but one DecNef participant. To investigate whether this bias, referred to hereafter as color-confidence bias, underlies the relationship between high confidence induction and false color perception during DecNef we performed a median split on DecNef study participants according to the difference in their mean confidence judgements for green versus red stimuli (Figure 7a). Replotting DecNef color induction likelihoods (Figure 4a) after median splitting suggested that DecNef color induction was not affected by color-confidence bias (Figure 7b).

**Figure 7.**
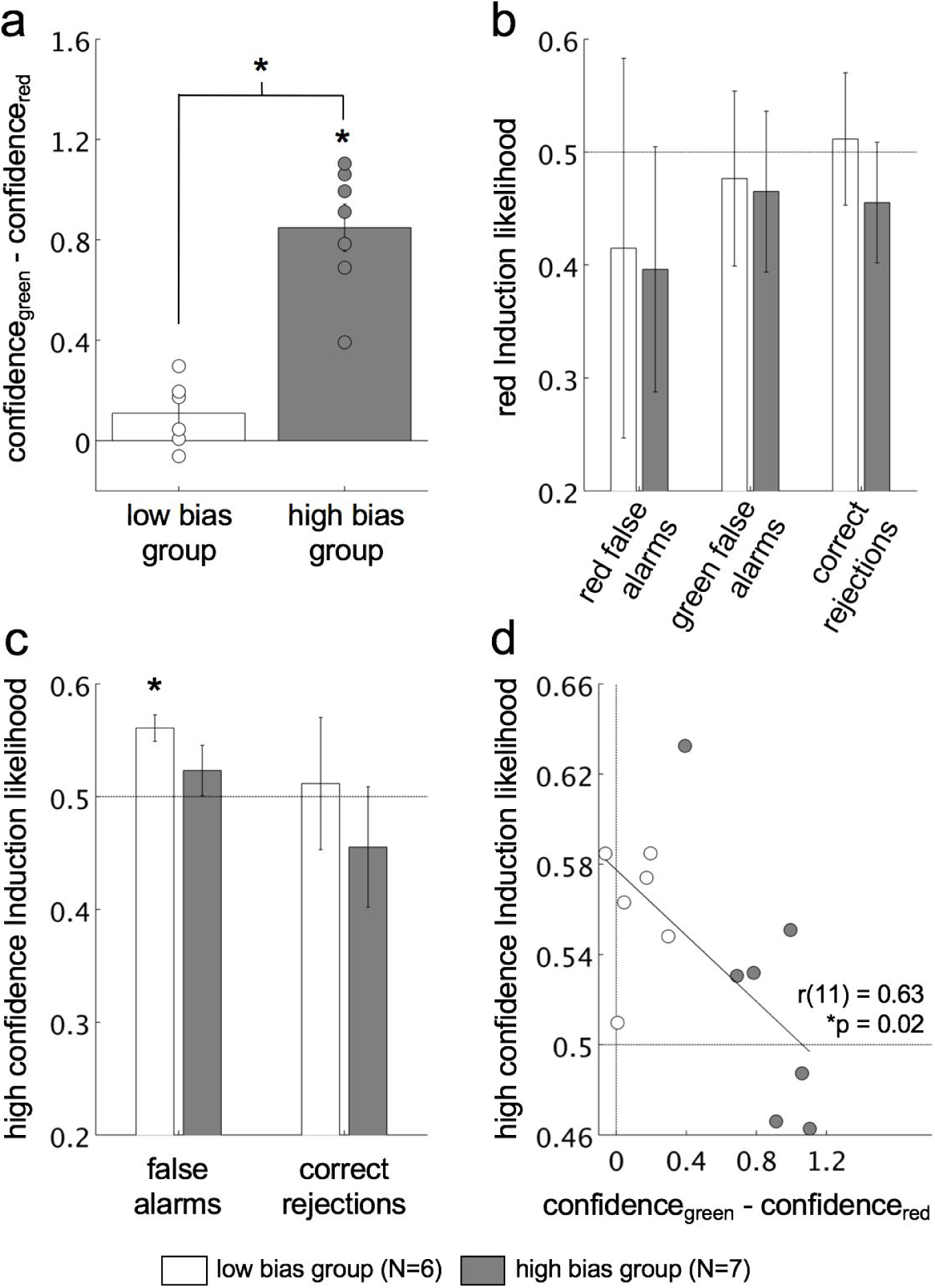
Median split analyses suggest that the association between high confidence induction and false alarms is not mediated by a bias for higher confidence in green decoder construction stimuli. **a)** A median split was conducted on the difference in mean confidence ratings for green stimuli and red stimuli. Because mean confidence ratings for red stimuli were subtracted from those for green stimuli, positive values suggest that, on average, subjects had higher confidence in green stimuli than red stimuli. As expected, the median split resulted in a significant difference in bias between the low bias group (white bar) and high bias group (gray bar) [t(11) = −6.47, p < 0.001, CI = (−0.99,-0.49), two-tailed, two-sample t-test]. Importantly, a one-tailed, one-sample t-test suggests that bias in the low group is not significantly different from zero at alpha = 0.05 [t(5) = 1.99, p = 0.052, CI = (−0.001,+inf)], although the low p-value suggests a trend in this direction. **b)** Red induction likelihoods for false alarm and correct rejection DecNef runs after median splitting on color-confidence bias. Median splitting showed no effect of color-confidence bias on red induction likelihoods during either false alarm or correction rejection runs. **c)** High confidence induction likelihoods for false alarm and correct rejection runs after median splitting on color-confidence bias. Induction likelihoods for participants in the low color-confidence bias group were significantly above chance (0.50) during false alarm runs [t(5) = 5.20, p < 0.01, CI = (0.53,0.59), two-tailed, one-sample t-test] but not during correct rejection runs. High confidence induction likelihoods were not significantly different from chance for either false alarm or correct rejection runs for participants in the high color-confidence bias group. **d)** Extent of color-confidence bias is inversely correlated with high confidence induction likelihoods during false alarm DecNef runs [r(11) = 0.63, p = 0.02].

To specifically test whether our main finding that high confidence induction likelihoods were higher during false alarm runs was affected by color-confidence bias we ran a mixed model ANOVA on high confidence induction likelihoods during DecNef with the within-subject factor run type (2 levels: false alarm runs, correct rejection runs) and the between-subjects factor color-confidence bias (2 levels: low, high). The ANOVA showed no main effect of either run type [F(1,11) = 2.62, p = 0.13] or color-confidence bias [F(1,11) = 1.01, p = 0.34] and no significant interaction [F(1,11) = 0.07, p = 0.80]. However, post-hoc Bonferroni corrected (α_corrected_ = 0.0125) two-tailed one-sample t-tests for each of each combination of bias group and run type showed that high confidence induction likelihoods were significantly above chance (0.50) in the low color-confidence bias group during false alarm runs [t(5) = 5.20, p < 0.01, CI = (0.53,0.59)], while high confidence induction likelihoods in each of the other three groups were not significantly different from chance [high bias, false alarm runs: t(6) = 5.20, p = 0.34, CI = (0.47, 0.58); low bias, correct rejection runs: t(5) = 0.20, p = 0.85, CI = (0.36, 0.66); high bias, correct rejection runs: t(6) = −0.84, p = 0.43, CI = (0.32, 0.59); Figure 7c]. Furthermore, there was a negative correlation between the extent of color-confidence bias and high confidence induction likelihoods during false alarm runs (R^2^ = 0.40, p = 0.02; Figure 7d). Taken together, these results suggest that color-confidence bias did not underlie the high high confidence induction likelihoods observed during false alarms DecNef runs. Conversely, the median split analysis suggests that this effect was most strongly driven by the study participants who showed the smallest color-confidence bias.

### Debriefing Questions

Induction strategies during neurofeedback ranged from actively trying to think nothing to imagining the induction stimulus being the top of a barbecue, on which meats were being grilled (Table 2). During the final debriefing session following the red/green color discrimination task on Day 7, eight out of 15 DecNef participants (53.3%) indicated on the forced-choice question that they thought they had been receiving sham neurofeedback. Further, only five out 15 DecNef participants (33.3%) responded that, assuming they had been receiving real neurofeedback, they were rewarded for activating a pattern of brain activity corresponding to red perception. These results suggest that participants were unaware of the true targets of neurofeedback.

**Table 2.** Examples of DecNef induction strategies. Participants were asked what strategies they employed during the DecNef task at the end of each neurofeedback session.

| Participant No. | Neurofeedback Strategies |
| --- | --- |
| 1 | imagined live rock music, playing sports, doing multiplication problems, a menu, working their old restaurant serving job, being a customer being served by their old self at that restaurant |
| 2 | imagined being happy, angry, sad, joyous; relaxed and mind wandered; counted prime numbers; imagined singing with friends; actively tried to think nothing |
| 3 | imagined vivid colors: sea and sky, green peppers, carrots; imagined playing soccer, basketball, baseball; imagined music and playing music, Monet paintings, puppies |
| 4 | imagined eating food -- it's appearance, texture, and taste; imagined chatting with friends, telling stories, the faces of people they'd seen the previous day, being rich, being an animal, getting hair done at a salon, getting married |
| 5 | imagined friends' faces after a car accident that they were in; tried to make themselves feel sad; imagined traveling; imagined painful things like running a marathon when you don't want to; imagined shouting angrily, singing; tried viewing the grating as clearly as possible; played word games |
| 6 | imagined what to eat for lunch; imagined a red grating (only 2 consecutive trials - the first made the feedback circle relatively large, the second did not); imagined the induction grating being the top of a barbecue on which they were grilling meats; imagined a green grating; imagined the induction grating being the bars of a jail cell, behind which was a famous baseball player who had been suspended for taking performance enhancing drugs; tried to empathize with that baseball player; wondered, "what is the meaning of this experiment"; imagined being in Las Vegas, viewing scenery through a moving train, the Grand Canyon, barbecue sauce, salt and pepper, what it would be like to run a business |
| 7 | imagined foreigners traveling through Japan; integrating equations in their mind; imagined being a train conductor, playing guitar, reading a music score, train station names, the story of a book they recently read, drawing difficult Hanzi; tried thinking about nothing; counted the seconds while the induction stimulus was on the screen; wished that the circle would get bigger |
| 8 | mental arithmetic, imagined music, singing, playing tennis and soccer, what that night's dinner would be, events that happened earlier that day; thought about their own research questions |
| 9 | tried to relax and think about nothing; imagined what it feels like to be in front of others at a swimming pool; imagined cutting and peeling the vertical bars of the grating, singing, walking on the black bars of the induction grating; tried to distribute attention widely across entire visual field |
| 10 | imagined past houses and school experiences, playing tennis, swimming, recent studies in telecommunications, the black disc around the fixation circle being blue, the fixation circle growing in size; focused on noise in the induction grating; imagined throwing a ball or shooting arrows at the fixation point |
| 11 | mental arithmetic: addition, multiplication, factoring; imagined running, doing track and field events; mind wandered; thought about how they could not control the size of the feedback circle; imagined the induction grating being green |
| 12 | mental arithmetic: addition, subtraction, multiplication, division; imagined writing sentences, scenery near their house, and playing soccer and baseball; |
| 13 | imagined friends' faces, feelings of depression, playing soft tennis, the smell of barbecued meat, vegetables, using non-dominant hand to pick up a bean with chopsticks; remembered getting an award and being praised at work, being afraid of high school tennis coaches, going on secret dates in high school |
| 14 | imagined singing, a woman dressed as a man singing, the shape of a train, potential names for a future child, a package of red apples, pictures of spouse, memories with spouse, cartoon characters, what they ate for lunch; focused on the induction grating; thought about what to make for dinner |
| 15 | imagined biking, the induction grating being a watery sphere, the induction grating being an archery target, the induction grating being green, scenes with blue skies and green lawns, friends' faces |

## Discussion

Using a DecNef paradigm that targeted activation of multivariate decoded patterns for color in visual cortex and perceptual confidence in frontoparietal cortex, we found that participants were more likely to activate patterns for high confidence during fMRI runs in which they also falsely perceived color. Activation of decoded patterns for color in visual areas, on the other hand, was not associated with false color perception.

### The relationship between confidence, consciousness, and prefrontal activity

The present data suggest that decoded patterns for perceptual confidence may be critically related to conscious visual perception. This provides support for confidence judgments in the extant debate about which subjective measure is ideal for measuring conscious awareness (Overgaard et al. 2010; Rosenthal 2018). Optimization of subjective measures is critical for the study of consciousness; the measure one selects can make the difference in whether or not a priming effect is considered to be truly subliminal (Wierzchoń et al. 2012) or whether above-chance orientation discrimination sensitivity is considered a case of Type 1 or Type 2 blindsight (Rausch and Zehetleitner 2016). In each of these cases, confidence judgments were found to be the most exhaustive and conservative measure of conscious awareness. While the current results do not rule out the possibility that decoded patterns for other subjective measures like visibility judgments might show a similar association with conscious perception in the same DecNef paradigm, they are informative nonetheless in providing at least a partial neural basis for the link between confidence judgments and consciousness.

These results also shed light on the current debate about whether prefrontal cortex is part of the core neural basis of consciousness (Melanie Boly et al. 2017; Odegaard, Knight, and Lau 2017). Given that three out of the four confidence ROIs were in prefrontal cortex, the observed association between confidence induction and conscious perception suggests that prefrontal cortex is critically involved in consciousness. Indeed, comparing high confidence induction under false alarm versus correct rejection runs in the individual frontoparietal ROIs as well as in a collective prefrontal ROI suggests that the main effect of high confidence induction under false alarms is driven by prefrontal activity (Figure 5a). Also supporting the idea that PFC is uniquely involved in the generation of conscious percepts are the result that the decoded patterns for color in visual cortex were not associated with false color perception (Figure 4b) and the relatively small extent to which activity in prefrontal ROIs could be predicted from activity in visual ROIs found in the information leak analysis (Figure 5b). Additionally, the fact that average high confidence induction likelihoods in confidence ROIs during neurofeedback were positively correlated with average confidence judgments (Figure 4c) supports the notion that the output of the confidence decoder was meaningfully related to perception.

Importantly, given that participants did not learn to activate the targeted induction patterns over time, we assume that the activation of decoded high confidence patterns during neurofeedback occurred spontaneously. The association between high confidence induction and false color perception therefore suggests an important role for ongoing spontaneous frontoparietal activity in determining the contents of conscious perception. This is in line with previous studies showing that perception is affected by spontaneous fluctuations in both sensory areas (Hesselmann et al. 2008, 2010; Sadaghiani, Hesselmann, and Kleinschmidt 2009; Samaha and Postle 2015; Samaha, Lemi, and Postle 2017) and frontoparietal areas (M. Boly et al. 2007; Rahnev et al. 2012; Sadaghiani, Hesselmann, and Kleinschmidt 2009).

### The effect of neurofeedback on color discrimination

The finding that participants’ red/green discrimination psychometric functions shifted in the direction of a higher overall proportion of red responses is consistent with a previous DecNef study targeting color representations in visual cortex (Amano et al. 2016). A potentially important difference, however, is that in the previous study it was reported that participants successfully learned to activate the targeted decoded color patterns more frequently over time through DecNef training, whereas in the current study such learning did not occur. One possible explanation for this difference is that despite the lack of learning in the current experiment, an association still formed between the induction stimulus, which was matched in its achromatic parameters to the target stimuli in the Day 3 and Day 7 psychophysics tasks, and spontaneous induction of decoded patterns for redness.

An alternative explanation is that the observed shift in psychometric functions in the current study was due to an initial Day 3 bias towards choosing green being minimized over time via non-DecNef-related perceptual learning. Future studies should investigate this by reducing such initial biases through longer training periods and more extensive stimulus titration based not only on task performance (as was the case here) but also on response bias measures like the signal detection theoretic criterion (Green DM 1966; MacMillan and Creelman 2004). Further, while here we rewarded all participants for activating the decoded pattern for red in line with Amano et al. (2016), it would be informative for future investigation to counter-balance by rewarding a second group of participants for activating the decoded pattern for green. A shift of the psychometric function in the opposite direction for the green group would provide strong evidence for an influence of DecNef on red/green color discrimination biases.

### Limitations & Future Directions

A limitation of the current study is the intermittent nature (i.e., run-by-run) of the psychophysical data collected during DecNef. Trial-by-trial measures of color perception would provide considerably greater power for evaluating the relationship between confidence and conscious awareness. Run-by-run psychophysical measures are not without precedent (Cheesman and Merikle 1984), and they were selected here as a means of reducing trial times. This was intended to facilitate participants learning to induce the targeted color and confidence patterns. However, given the lack of such learning observed here, it may be optimal for future studies to prioritize the greater power and signal-to-noise ratio afforded by trial-by-trial perceptual judgments.

It remains an open question, however, what prevented participants from learning to activate the targeted color and confidence patterns in the DecNef task. As such an effect would allow for the investigation of a causal relationship between perceptual confidence and consciousness, this is an important issue for future studies to investigate. The present DecNef study was the first to use two categorically different perceptual targets (color and confidence); one possibility, therefore, is that the combination of these patterns is too complex for participants to learn to generate consistently. Another related possibility is that the neurofeedback procedure was simply spread across too many decoders (one for color and four for confidence). These issues may have also been responsible for the failure of confidence neurofeedback to modulate confidence judgements as previously reported (Cortese et al. 2016).

Future studies should investigate the limits of what human participants can learn to regulate via DecNef in terms of both the number and distribution of decoders throughout the brain, and the complexity of the perceptual content being targeted. For example, it is an open question whether confidence DecNef would benefit from using a single decoder spanning the four frontoparietal ROIs used here, as might be suggested by successful approaches using whole brain decoders (deBettencourt et al. 2015). It would also be informative to repeat the current study with confidence DecNef alone, and to limit the neurofeedback to only prefrontal ROIs (Figure 5a). If learning occurred in either of these modified contexts, or some combination of them, then it would suggest that the difference in learning indeed stems from the difference in either the number of decoders or the categorical complexity of the targets of neurofeedback.

It is also possible that better decoding accuracy would have led to induction learning. One recent study showed that decoding accuracies could be significantly improved by integrating fMRI hyperalignment (Haxby et al. 2011) into the DecNef procedure (Taschereau-Dumouchel et al. 2018). Hyperalignment achieves this by taking advantage of shared high-dimensional patterns in representational content between subjects. Another study showed that offline simulations can be used to estimate optimal parameters for experiment timing and real-time fMRI preprocessing, which can lead to greater decoding accuracy and neurofeedback performance (Oblak, Sulzer, and Lewis-Peacock 2019). Similar approaches should be applied going forward to optimize the efficacy of DecNef tasks, thereby giving us greater power to investigate the relationship between confidence and consciousness.

## Acknowledgements

We thank K. Nakamura for her help with scheduling and conducting the experiment and Y. Shimada and A. Nishikido for operating the fMRI scanner. The study was conducted in the ImPACT Program of Council for Science, Technology and Innovation (Cabinet Office, Government of Japan).

## Data Availability

The data that support the findings of this study are available from the corresponding author upon reasonable request.

